## Supplemental Figures for "Predicting cell cycle stage from 3D single-cell nuclear-stained images"

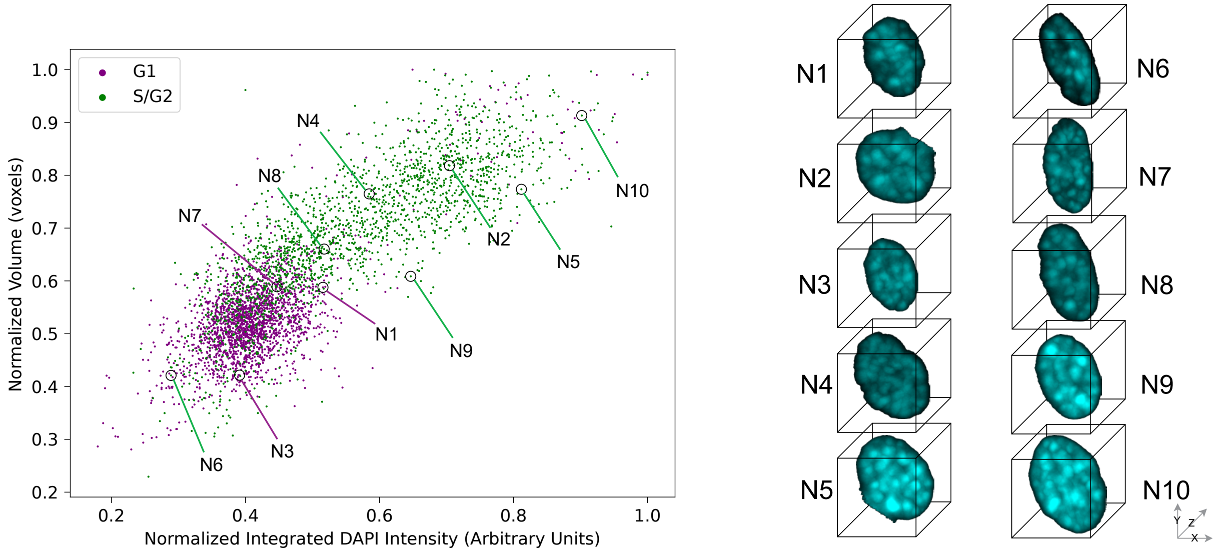


Figure S1: **Scatter plots of epifluorescence image dataset with top 10 representative cell images.**

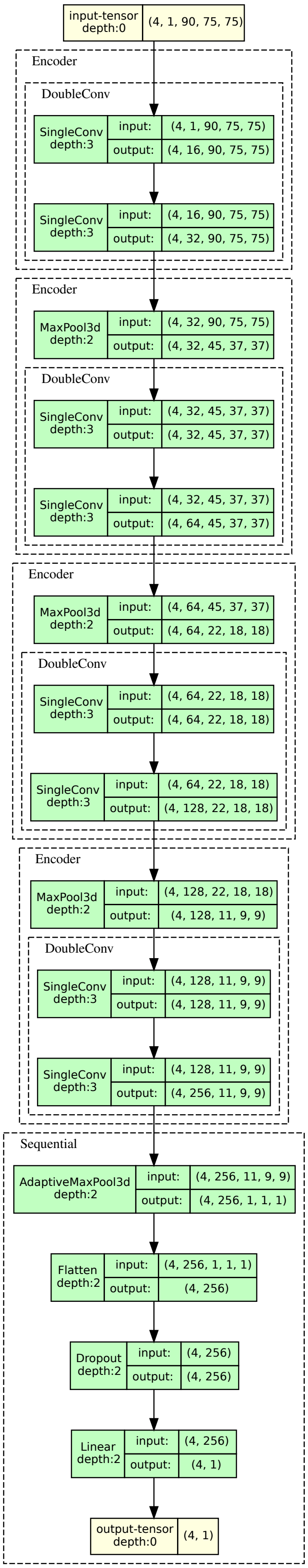


Figure S2: **Architecture details of CellCycleNet with a batch of 4 images.**

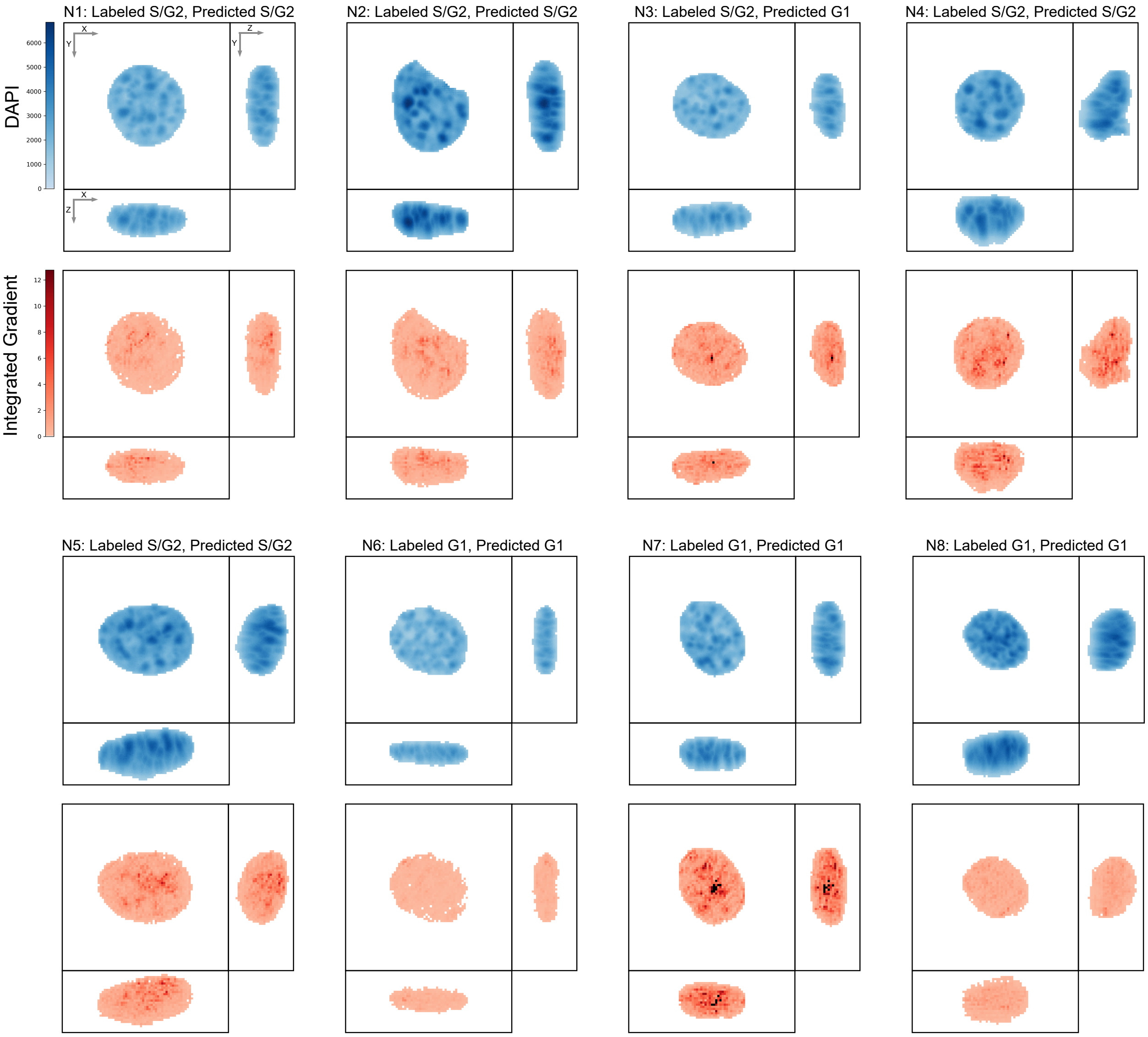


Figure S3: **Integrated gradients of CellCyleNet for top 8 representative cells of confocal dataset.**

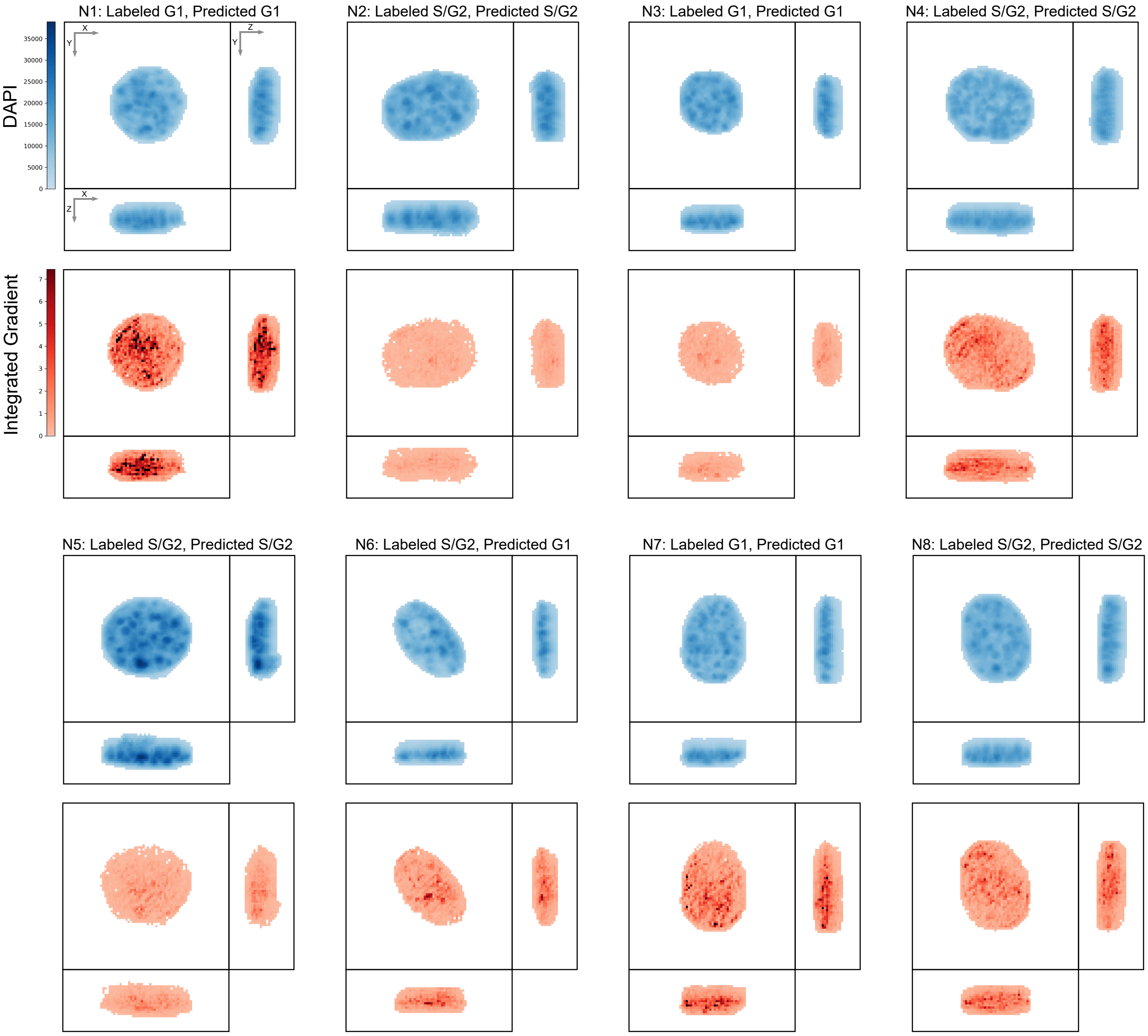


Figure S4: **Integrated gradients of CellCyleNet for top 8 representative cells of the epifluorescence dataset.**
